# Truly dorsal: nonlinear integration of motion signals is required to account for the responses of pattern cells in rat visual cortex

**DOI:** 10.1101/2023.02.17.528967

**Authors:** Giulio Matteucci, Rosilari Bellacosa Marotti, Benedetta Zattera, Davide Zoccolan

## Abstract

The defining feature of advanced motion processing in the primate dorsal stream is the existence of pattern cells – specialized cortical neurons that integrate local motion signals into pattern-invariant representations of global direction. Pattern cells have also been reported in rodent visual cortex, but it is unknown whether the tuning of these neurons results from truly integrative, nonlinear mechanisms or trivially arises from linear receptive fields (RFs) with a peculiar geometry. Here we show that pattern cells in rat visual cortical areas V1 and LM process motion direction in a way that cannot be explained by the linear spatiotemporal structure of their RFs. Instead, their tuning properties are consistent with those of units in a state-of-the-art neural network model of the dorsal stream. This suggests that similar cortical processes underly motion representation in primates and rodents. The latter could thus serve as powerful model systems to unravel the underlying circuit-level mechanisms.

## Introduction

Perceiving the velocity (i.e., motion direction and speed) of visual objects is critical to interact effectively with the environment. From a computational point of view, primary visual cortex (V1) can be thought as a bank of local moving-edge detectors. Upstream sensory areas face the nontrivial challenge of extracting object motion direction from the V1 representation. The output of a single, localized edge-detector is, in fact, intrinsically ambiguous, since it reflects the projection of the global velocity vector onto the direction that is orthogonal to the orientation detected by the unit. Any information regarding the component of object motion that is parallel to such orientation is lost. Thus, when considered in isolation, the output of each of these edge detectors is compatible with infinite combinations of global directions and speeds and is therefore insufficient to fully specify the velocity of the underlying object. Only by combining (i.e., integrating) multiple local direction signals of this kind, global object direction and speed can be fully determined. This ambiguity is known in the neuroscientific literature as the “aperture problem” (Fennema and Thompson, 1979; Marr, 2010; Wuerger et al., 1996). Psychophysically, it can be appreciated by the fact that observers looking at a drifting object through a small aperture will perceive the edge seen through the aperture as always drifting in the perpendicular direction to the edge itself, irrespectively of the global direction of the object behind the aperture (Wuerger et al., 1996). If not handled properly by the visual system, the aperture problem would lead to illusory and inaccurate motion measurements (see Supplementary Video 1).

In the brain of primates, motion integration is known to be achieved by pattern cells, which are abundant in monkey dorsal stream areas such as MT (Khawaja et al., 2013, 2009; Movshon et al., 1985; Movshon and Newsome, 1996; Rodman and Albright, 1989; Rust et al., 2006; Smith et al., 2005; Solomon et al., 2011) and MST (Khawaja et al., 2013, 2009). The complementary class of cells is known as component cells – neurons that, behaving more like the local moving-edge detectors described above, are sensitive to the aperture problem. This class of neurons has been reported to be predominant in V1 (Khawaja et al., 2013, 2009; Movshon et al., 1985; Movshon and Newsome, 1996) and widespread across multiple areas of the monkey visual cortex.

In rodents, only a handful of studies have investigated the distribution of pattern and component cells in mouse V1 (Muir et al., 2015; Palagina et al., 2017) and bordering high-order visual areas (Juavinett and Callaway, 2015). Such studies yielded contrasting results about the presence of pattern cells in V1: two of them reported a small but consistent fraction of pattern units in this area (Muir et al., 2015; Palagina et al., 2017), while another did not find any, reporting instead their presence in high-order areas LM and RL (Juavinett and Callaway, 2015).

Despite these contrasting findings on the neurophysiological front, recent work has provided compelling evidence about the causal involvement of V1 in mediating discrimination of motion direction of random dot fields in mice (Marques et al., 2018). Moreover, in a previous study, we have shown that rats can spontaneously perceive global motion direction of drifting plaids – i.e., composite patterns, made of two superimposed gratings drifting along different directions, which are typically used to distinguish pattern from component cells (Matteucci et al., 2021). In that study, rats trained to discriminate plaids drifting along opposite directions successfully (and spontaneously) generalized their discrimination to drifting gratings, thus displaying an ability consistent with the existence of pattern cells. However, rats trained with drifting gratings did not generalize to drifting plaids. This unexpected finding prompted some computational modeling work suggesting two alternate scenarios about the properties of motion representations in rat visual cortex: one, in which decision neurons learn to rely preferentially on the output of pattern cells (when rats are trained with plaids), under the hypothesis that component cells are more strongly affected by cross-orientation suppression than pattern cells; and an alternative scenario, where decision neurons are “hard wired” to read out the output of pattern cells, which are as strongly affected by cross-orientation suppression as component cells. Which of these scenarios account for the behavioral findings in (Matteucci et al., 2021) critically depends on the relative impact of cross-orientation suppression on rat component and pattern cells, which is currently unknown.

More in general, the limited evidence about a hierarchical growth of pattern cells along the mouse putative dorsal stream, their overall paucity, and the fact that they have been found as early as in V1 raise the question of whether these neurons truly perform those nonlinear, integrative computations that are typical of primate pattern cells. In fact, as originally proposed by (Tinsley et al., 2003), a unit with a Gabor-like, linear receptive field (RF) can be tuned to the global motion direction of a plaid, thus displaying a pattern-like behavior, if the patches of local luminance in the plaid tightly overlap with the excitatory/inhibitory subfields of the cell’s RF (see cartoon in Fig. 1C). The shorter and wider (i.e., the “blobbier”) the RF subfields are (compare cartoons in Fig. 1B and C), the more pattern-like the direction tuning of the cell will look like. Following (Tinsley et al., 2003), we quantitively probed this scenario by simulating edge-detector units using Gabor filters with different aspect ratios and by measuring their responses to gratings and plaids (with a 120° cross-angle) drifting along 12 equi-spaced drift directions (Fig. 1A). Filters with high aspect ratio (top row) produced sharp tuning curves for both the grating (dotted line) and plaid (solid line) stimuli. Moreover, the tuning curve for the plaids peaked at +/- 60° (i.e., half plaid cross-angle) from the preferred direction of the gratings – the signature property of component cells. For filters with an intermediate aspect ratio (middle row), the tuning curves became broader and the two peaks of the curve obtained for the plaids partially overlapped, displaying a tendency to merge. Finally, for very low aspect ratios (bottom row), the further broadening of the tuning curves led to a complete merge of the two peaks of the curve obtained for the plaids into a single peak – thus yielding curves centered on the same (global) direction for both plaids and gratings. This shows how the defining property of pattern cells (i.e., similar tuning curves for gratings and plaids) could arise from purely linear spatial filters by virtue of their geometry (Fig. 1B-C).

**Fig. 1.**
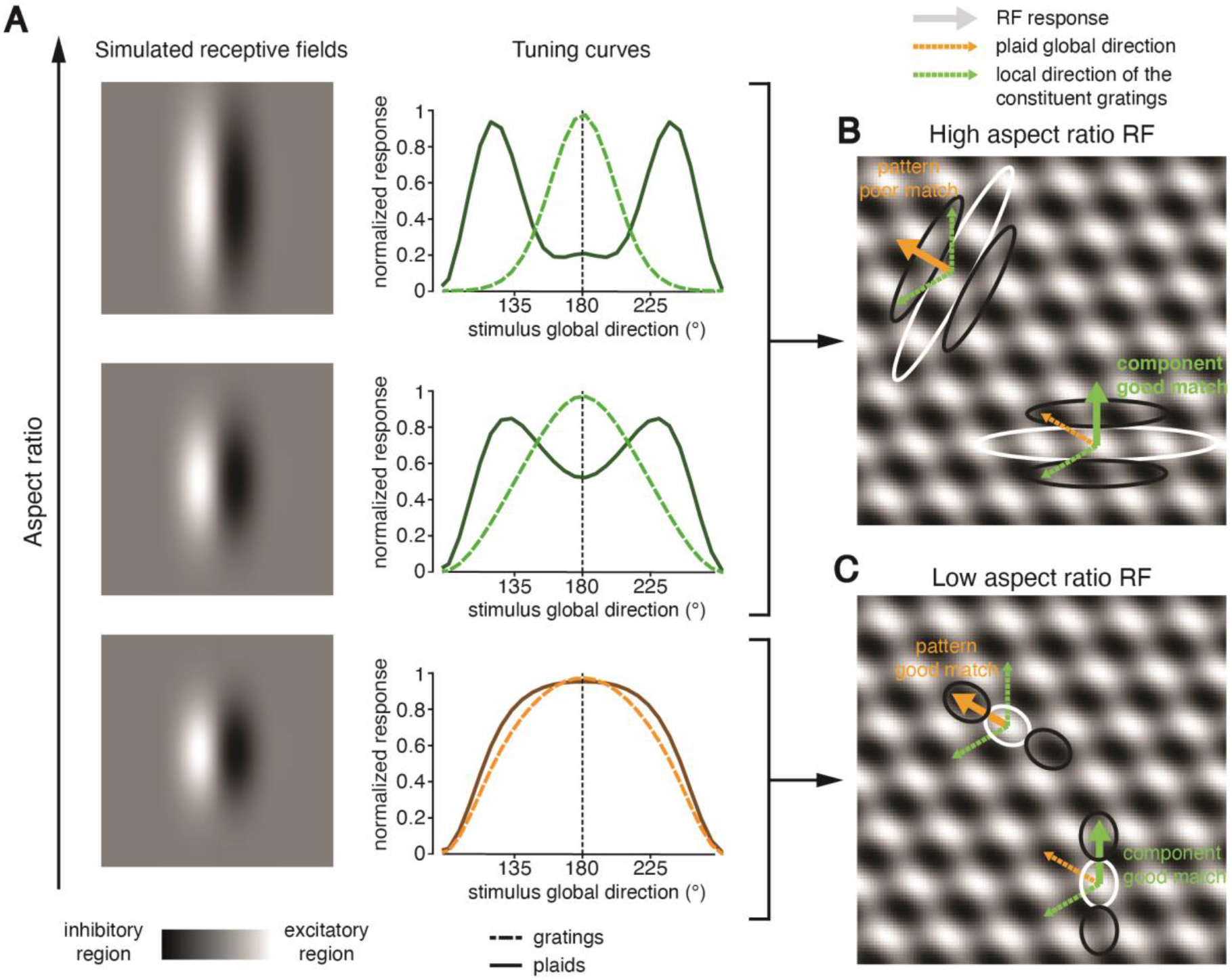
Illustration of how “blobby”, linear receptive fields can produce tuning consistent with the behavior of pattern cells. (**A**) Left: spatial structure of the linear filters used to simulate receptive field (RFs) with progressively lower aspect ratio (from top to bottom) using Gabor functions. Right: tuning curves showing the selectivity for grating and plaid direction (respectively, dashed and solid lines) of the linear filters shown on the left. The lower is the aspect ratio of the RFs, the broader is the tuning of the simulated units and the stronger is the tendency of the curves obtained for gratings and plaids to overlap (i.e., to display the tuning expected for pattern cells). (**B-C**) Graphical illustration of how the aspect ratio of linear, Gabor-like RFs (as those simulated in A) can determine the extent to which the selectivity of a unit is consistent with the tuning of either a component or a pattern cell (adapted from (Tinsley et al., 2003)). Panel B depicts a case where an elongated RF with high aspect ratio poorly matches the local contrast features of a plaid that drifts at 135° (orange arrow), being made of two superimposed gratings, one drifting vertically (at 90°) and the other one drifting at 210° (green arrows). When the RF is oriented in such a way to be orthogonal to the local direction of one of the constituent gratings (see example Gabor on the bottom/right), there will be times at which the average luminance falling within the subunits of the RF matches their polarity, thus producing a strong response. If instead the RF is orthogonal to the global direction of the plaid (see example Gabor on the top/left), the average luminance falling within each subunit of the RF is approximately mid-gray, thus producing no response. As a result, the unit can detect and signal the local direction of the component (thick green arrow) but not the global direction of the plaid (thick orange arrow). Panel C depicts a very different case, where a “blobby” RF with low aspect ratio matches well the local contrast features of the plaid (same stimulus as in B). No matter whether the orientation of the RF is orthogonal to the local direction of one of the constituent gratings (see example Gabor on the bottom/right) or to the global direction of the plaid (see example Gabor on the top/left), there will be times at which the average luminance falling within the subunits of the RF matches their polarity, thus producing strong responses. This will allow the unit to respond in a similar way to gratings and plaids drifting along the same direction, yielding the pattern cell tuning shown in A (bottom plot).

As already pointed out by (Palagina et al., 2017), the mechanism illustrated above implies the possibility that pattern selectivity observed in rodent visual cortex could simply be the result of linear RFs with low aspect ratio and broad direction tuning. This scenario is consistent with the fact that, compared to mammals with higher visual acuity (De Valois et al., 1982b, 1982a; Gizzi et al., 1990; Ringach et al., 2002), rodent visual neurons have indeed broader tuning curves, preferences for lower spatial frequencies (Girman et al., 1999; Marshel et al., 2011; Niell and Stryker, 2008), and RFs with a lower aspect ratio (Matteucci et al., 2019; Niell and Stryker, 2008). In addition, although (Palagina et al., 2017) reported no significant difference of average tuning broadness between pattern and component cells in mouse V1, they found that pattern responses are not cross-angle invariant (i.e., they change their pattern/component behavior depending on the angle between the two component gratings forming the plaid) – a signature of a possible dependence from RF geometry.

In the light of these considerations, when investigating rodent visual cortex, the risk of misclassifying linear, non-integrative units (that should be properly considered as broadly tuned component cells) as pattern cells cannot be overlooked. This could explain pattern selectivity observed in rodents without invoking nonlinear hierarchical mechanisms similar to those thought to be at work in primate cortex (Rust et al., 2006; Simoncelli and Heeger, 1998).

The most direct way to test for this possibility is to reconstruct the linear receptive fields of putative pattern cells and try to predict their plaid and grating responses on that basis. If the geometry of their RFs is the cause of their pattern-like tuning, the responses predicted by linear RFs should still be pattern-like. Conversely, if nonlinear, truly integrative mechanisms are at work, linear RFs should fail to produce pattern-like responses.

Our study was designed to: i) address this question about the nature of rodent pattern and component cells; ii) assess their sensitivity to cross-orientation suppression, so as to test the two alternative hypotheses put forward in (Matteucci et al., 2021); and iii) measure their relative abundance across rat V1 and the two extrastriate areas (LM and RL), which, in mice, have been reported to contain the larger fraction of pattern cells (Juavinett and Callaway, 2015). Finally, the entire data processing pipeline applied to the neuronal data was also employed to characterize the RF structure and tuning properties of the units of a state-of-the-art neural network model of the monkey dorsal stream (Mineault et al., 2021). This allowed a quantitative assessment of the level of sophistication of motion processing by rat visual cortical neurons and an indirect comparison with primate visual cortex.

## Results

We performed extracellular recordings from primary visual (V1), lateromedial (LM) and rostrolateral (RL) cortex of anesthetized rats. The animals were presented with a battery of stimuli including: 1) gratings and plaids (with a 120° cross-angle), drifting along 12 equi-spaced directions (from 0° to 330°) and presented at two different spatial and temporal frequencies (SFs = 0.02 and 0.04; TFs = 4 and 6); and 2) spatiotemporally correlated noise movies (see Materials and methods). Grating and plaid responses were used to compute direction tuning curves and classify the recorded single units as pattern or component cells, based on the standard approach developed in cat and monkey studies (Gizzi et al., 1990; Movshon et al., 1985; Rodman and Albright, 1989; Scannell et al., 1996; Solomon et al., 2011), and also used in previous rodent studies (Juavinett and Callaway, 2015; Muir et al., 2015; Palagina et al., 2017). Noise movies were used to obtain a linear estimate of the spatiotemporal RF of each neuron (i.e., to find the best linear filter that approximated the stimulus-response function) by using the Spike-Triggered Average (STA) technique (Aljadeff et al., 2016; Schwartz et al., 2006; Sharpee, 2013).

We recorded a total of 447, 367 and 412 well-isolated single units from, respectively, V1, LM and RL. Among these neurons, 258 units in V1, 187 in LM and 184 in RL were significantly responsive to gratings or plaids (see Material and methods) and were therefore included in the analyses described below. Following previous studies (Gizzi et al., 1990; Movshon et al., 1985; Rodman and Albright, 1989; Scannell et al., 1996; Solomon et al., 2011), “patternness” and “componentness” were quantified by computing the Fisher-transformed partial correlation between the observed responses to the plaids and the predicted responses to the same stimuli, as inferred from the observed responses to the gratings, assuming either an ideal pattern or component selectivity (these correlations are referred to as Z_p_ and Z_c_ respectively; see Materials and methods for a definition). Direction selective neurons with Z_p_ significantly higher than 0 and larger than Z_c_ were classified as pattern cells; vice versa, direction selective units with Z_c_ significantly higher than 0 and larger than Z_p_ were classified as component cells (see Material and methods). Neurons that did not meet one of these requirements were labelled as unclassified.

Figure 2 shows the tuning of a few example neurons recorded from the three targeted areas and classified as either component or pattern cells. For each unit, the figure reports: (1) the normalized tuning curve as a function of the direction of the stimulus (either a grating, left, or a plaid, right), when presented at the most effective SF and TF (solid lines); (2) a sequence of STA images, computed at progressively larger time lags from the firing of an action potential, showing the spatiotemporal evolution of the RF structure; and (3) the tuning curves (dashed lines) predicted by a linear-nonlinear (LN) model of stimulus–response mapping using the spatiotemporal filter estimated via the STAs (Aljadeff et al., 2016; Schwartz et al., 2006; Sharpee, 2013). The figure also reports the values of the metrics used to quantify the response properties of the neurons, i.e.: 1) the “patternness” and “componentness” indexes for both the observed (Z_p_ and Z_c_) and predicted (Z_p_’ and Z_c_’) tuning curves; and 2) the contrast index (CI) used to quantify the sharpness of the STA images (Matteucci et al., 2019; Matteucci and Zoccolan, 2020).

**Figure 2.**
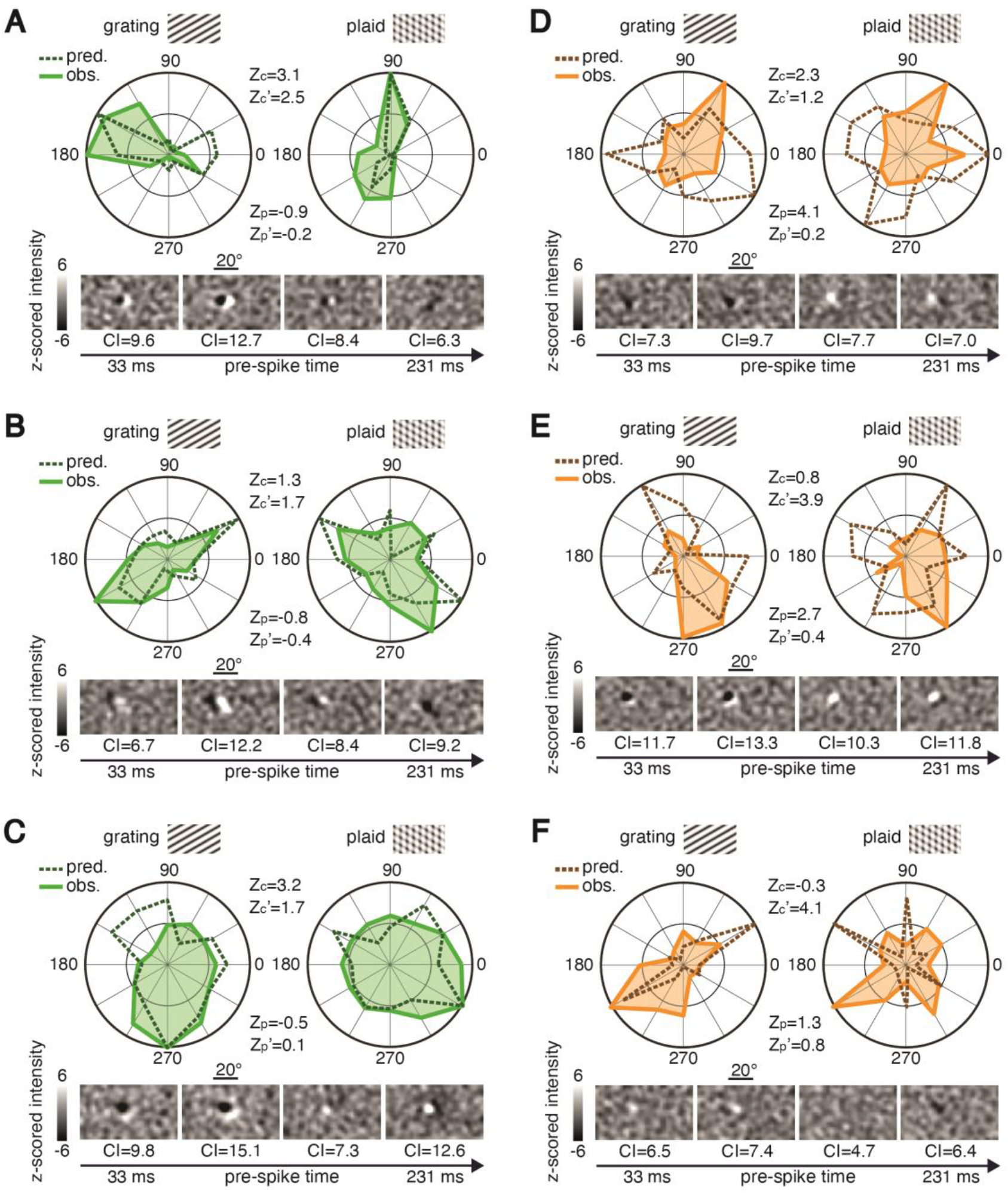
Examples of pattern and component cells recorded in areas V1, LM and RL. Each panel depicts the observed tuning curves (i.e., normalized average response for each stimulus direction) for gratings (left; solid line) and plaids (right; solid line), as well as the tuning curves predicted by a LN model based on the spatiotemporal filter estimated via STA (dashed lines). The pairs of values Z_c_ and Z_p_ and Z_c_’ and Z_p_’ are the component and pattern indexes computed for, respectively, the observed and predicted tuning curves. The temporal sequence of STA images at different time lags preceding spike generation is also shown (bottom), along with the CI values that quantify the sharpness of each STA filter. In every STA image, each pixel intensity value was independently z scored, based on the null distribution of STA values obtained through a permutation test (see Materials and Methods). A gray scale map was used to visualize the resulting z-scored values within the [-6, +6] range (see scalebars). (**A**) Sharply tuned component cell from V1, correctly predicted by the LN model as component. (**B**) Sharply tuned component cell from LM, correctly predicted by the LN model as component. (**C**) Broadly tuned component cell from V1, correctly predicted by the LN model as component. (**D**) Sharply tuned pattern cell from LM, incorrectly predicted by the LN model as unclassified. (**E-F**) Sharply tuned pattern cells from LM, incorrectly predicted by the LN model as component.

Figure 2A shows a typical V1 component cell displaying clear direction selectivity. When probed with the gratings, the unit’s direction tuning curve featured a well-defined peak in the 150°-180° range. By contrast, when measured with the plaids, the curve displayed two peaks at about ±60° with respect to the direction of the preferred grating. This inconsistency between the peak responses obtained for gratings and plaids is what made the unit a component cell, i.e., a poor global motion detector. In fact, the neuron was not sensitive to the actual direction of the plaid, but to the direction of its constituent gratings – only when the latter aligned with the preferred grating direction, the unit fired vigorously. The figure also shows how STA returned images with large CI values containing crisp, Gabor-like RFs made of two flanking lobes (one excitatory and one inhibitory), aligned along an axis that was orthogonal to the unit’s preferred orientation, and with a phase that gradually shifted over time, consistently with the strong direction selectivity of the neuron. The STA images yielded a good approximation of the unit’s spatiotemporal RF, as demonstrated by the close match between the measured tuning curves and those predicted by the LN model for both the grating and plaid responses (compare solid and dashed lines). As a result, Z_c_ was considerably larger than Z_p_ for both the observed and predicted tuning curves, yielding a classification of the cell as *component* in both cases. As similar behavior can be observed for two other example cells – one recorded in LM, having a sharp orientation tuning curve when tested with gratings (Fig. 2B), and another one, recorded in V1, with broader tuning (Fig. 2C). In both cases, the tuning curve obtained with the plaids featured two peaks, roughly at ±60° with respect to the direction of the preferred grating, and the spatiotemporal RF was well captured by STA, yielding, again, multi-lobed, Gabor-like filters (with large CI values) that accurately predicted grating and plaid responses via the LN model. As a result, Z_c_ was larger than Z_p_ for both the observed and predicted curves, indicating that both units were component cells and were correctly predicted as such by the LN model.

A different behavior can be observed for the three example LM neurons shown in Fig. 2D-F (orange curves). In all cases, the units were narrowly tuned for a specific grating direction, and such preference was maintained when tested with the plaids, yielding, for both stimulus classes, highly consistent, sharply tuned curves. This is the typical tuning expected for pattern cells, as confirmed by the larger magnitude of Z_p_, as compared to Z_c_. Interestingly, this behavior was not captured by the LN model. In the case of the first cell (Fig. 2D), STA returned a “blobby” RF with a main, dominant lobe and lower CI (on average, across frames), as compared to the component cells shown in Fig. 2A-C. As a result, the STA-based LN model only partially accounted for the tuning for grating direction and fully failed to predict the tuning for plaid direction (compare solid and dashed lines). In the case of the second cell (Fig. 2E), STA images with high contrast (CI) were obtained across almost the entire spatiotemporal evolution of the RF. The resulting LN model successfully predicted the tuning of the unit for grating orientation, although not for direction. Critically, however, the LN model failed to account for the tuning for plaid direction, yielding a predicted tuning curve, with peaks at about ±60° with respect to the direction of the preferred grating, consistent with the behavior of a component rather than a pattern cell. A similar behavior was observed for the third cell (Fig. 2F), whose RF, as recovered by STA, despite being poorly structured and with rather low CI, succeeded at roughly capturing the tuning of the unit for grating orientation (but, again, not for direction). However, as for the previous neuron, the LN model failed to predict the tuning for plaid direction, returning a curve that was inconsistent with the behavior of a pattern cell, having peaks at about ±60° with respect to the preferred grating direction. As a result, although the three neurons in Figs. 2D-F were classified as pattern cells, based on the observed responses to gratings and plaids, none of them retained such classification when their predicted responses via the LN model were considered – based on such predictions, the cell shown in Fig. 2D fell in the *unclassified* category, while the cells shown in Fig. 2E-F were classified as *component*.

### Incidence and susceptibility to cross-orientation suppression of component and pattern cells in cortical areas V1, LM and RL

The example neurons shown in Fig. 2 demonstrate that rat visual cortical areas do contain neurons that can robustly be classified as either pattern or component cells. Fig. 3A-C reports the incidence of these cell types in the investigated areas, by plotting, for each unit, the pair of Z_p_ and Z_c_ indexes (the anatomical location of the areas over the cortical surface is indicated in Fig. 3D). The boundaries in the figures indicate regions of the Z_p_/Z_c_ plane where one of the indexes is sufficiently larger than zero, as well as sufficiently larger than the other one, for a cell to be classified as either component (in green; bottom-right corner) or pattern (in orange; top-left corner). All other neurons were considered unclassified (in gray; central region). In addition, neurons were labeled according to whether they were sufficiently direction tuned (darker shades; DSI > 0.33) – an additional requirement that allows classifying a unit as a component (or pattern) direction selective cell (see Materials and methods).

**Figure 3.**
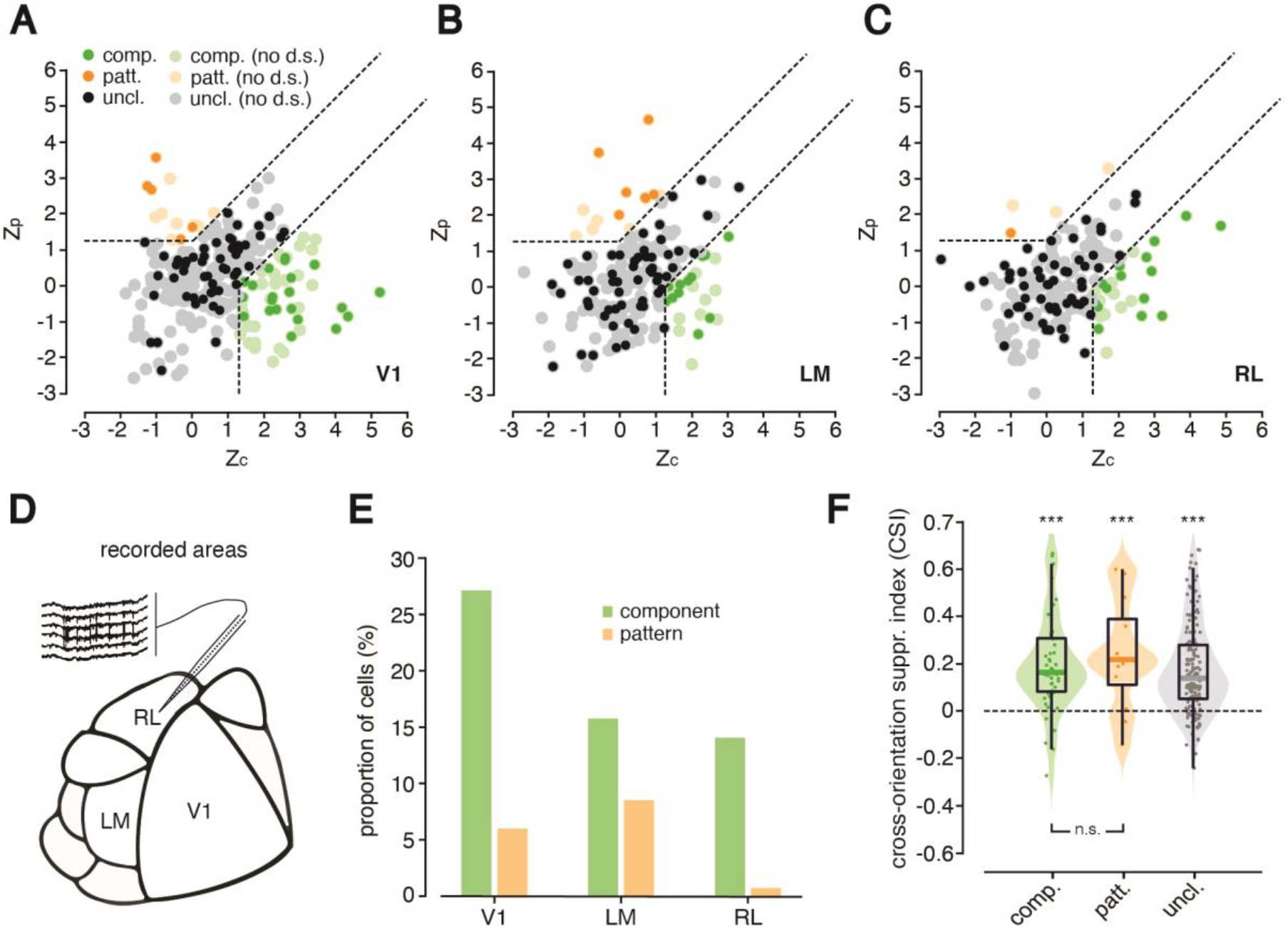
Distribution of pattern and component cells in areas V1, LM and RL. (**A-C**) Scatter plots showing the distributions of the pairs of observed Z_p_ and Z_c_ indexes for the cells recorded in the three targeted areas. Light-colored dots represent units not meeting the direction selectivity criterion (i.e., DSI > 0.33); dark-colored dots represent units meeting such criterion. Dashed lines are decision boundaries in the Z_p_/Z_c_ plane that distinguish regions where cells are labeled as component (in green; bottom-right corner), pattern (in orange; top-left corner) or unclassified (in gray; central area). (**D**) Schematic map of rat visual cortex showing the anatomical locations of V1, LM and RL. (**E**) Percentage of component cells (green bars) and pattern cells (orange bars) recorded in V1 (left), LM (center) and RL (right). (**F**) Distributions of cross-orientation-suppression index (CSI) values across area-pooled populations of component (left; green), pattern (center; orange) and unclassified (right; gray) units. All medians were significantly larger than zero (\*\*\**p* < 0.001; Wilcoxon test). The medians of the component and pattern cells’ pools were not statistically different from each other (*p* > 0.05; Wilcoxon test).

As shown in Fig. 3E, in agreement with (Palagina et al., 2017) and (Muir et al., 2015), but in contrast with (Juavinett and Callaway, 2015), our recordings yielded a sizable amount of pattern cells in V1 (6% of the responsive 78 units meeting the direction selectivity criterion). On the other hand, in qualitative agreement with the latter study, we found a significant fraction of pattern cells in LM (9% of the 76 units meeting the direction selectivity criterion). By contrast, we found in RL only a single pattern cell (i.e., about 1% of the 73 units meeting the direction selectivity criterion). The fraction of component cells in V1 (27% out of 78 units) was found to be in qualitative agreement with (Muir et al., 2015) and (Juavinett and Callaway, 2015), but larger than the one reported by (Palagina et al., 2017) (where it amounted to about half the figure we observed). On the other hand, whereas the proportion of component cells we observed in RL (21% out of 73 units) was similar to the one observed by (Juavinett and Callaway, 2015), the fraction in LM (16% out of 76 units) was roughly half of what they reported. Finally, consistently with previous rodent studies, the largest fraction of single units included in the analysis fell into the unclassified category (67% in V1, 76% in LM and 77% in RL).

Overall, these results confirm that, also in rats, as previously observed in mice, motion sensitive neurons exist that can be classified as either component or pattern cells, according to the criteria commonly adopted in the monkey literature. In addition, we observed a small increase in the proportion of pattern cells from V1 to LM, paralleled by a decrease in the proportion of component cells. Although such a tradeoff in the relative abundance of the two cell types did not reach statistical significance (χ^2^ test, *p* = 0.1870), we also observed a significant increase of the average value of the Pattern Index (PI = Z_p_-Z_c_, a commonly used scalar metric of “patternness) from V1 (PI = −1.64 +/- 0.31) to LM (PI = −0.57 +/- 0.40; unpaired t test p < 0.05). Taken together, these observations are suggestive of a shallow, hierarchical buildup of global motion detectors from V1 to LM.

Next, we checked the extent to which these cell types are affected by cross-orientation suppression, given the relevance of this property to probe alternative perceptual models of motion integration in rats (Matteucci et al., 2021). To measure this phenomenon, we computed a cross-orientation suppression index (CSI), which quantifies the amount of suppression (or facilitation) of neuronal firing when a unit is tested with its preferred plaid stimulus, as compared to its preferred grating stimulus. That is, the index is defined as the relative firing rate difference between peak grating and plaid responses (see Material and Methods): a positive value indicates cross-orientation suppression (i.e., larger peak response for gratings than for plaids), while a negative value indicates facilitation (i.e., larger peak response for plaids than for gratings). As shown in Fig. 3F, the CSI values for both component and pattern cells were shifted towards positive values (green and orange distributions), with the medians of the two populations being significantly higher than zero (*p* < 0.001; Wilcoxon test) but not statistically different from each other (*p* > 0.05; Wilcoxon test). The same trend was observed for the pool of unclassified units (gray distribution). Thus, for both component and pattern cells, the presence of multiple overlapping gratings leads to a suppression of the firing rate, as compared to the case where a single, isolated grating is used as a visual stimulus. The implication of this result for understanding the perceptual mechanisms underlying motion integration in rats are examined in the Discussion.

### Linear spatiotemporal receptive fields are sharper and more structured for component than for pattern cells

The main goal of our study, beside assessing whether pattern cells are present in rat visual cortex, was to understand whether the tuning of these units can properly be ascribed to nonlinear integrative mechanisms or, instead, is mainly the result of linear filtering via “blobby” RFs (see Introduction and Fig. 1). To address this question, we measured the time evolution of spatial RFs at ten progressively longer time lags from spike generation using STA analysis (see examples in Fig. 2). In this section, we test the hypothesis that STA better captures the RFs of component than pattern cells, as expected if the stimulus-response relationship was more nonlinear for the latter.

As illustrated by the example units of Fig. 2 and by the additional examples of Fig. 4A, component cells do appear to have RFs that are somewhat sharper (i.e., with larger contrast, as compared to the background noise) than those of pattern cells, and closer to Gabor functions containing at least two lobes. In Fig. 4, following an approach we already adopted in previous studies (Matteucci et al., 2019; Matteucci and Zoccolan, 2020), we statistically quantified this comparison, by plotting the distributions of: 1) the CI values obtained for the STA images in the two populations of component and pattern cells at all tested (ten) time lags from spike generation (Fig. 4B); 2) the goodness of fit (GOF) of all these STA images with Gabor functions (i.e., fraction of explained variance; Fig. 4C); and 3) the number of distinct, prominent lobes in the sharpest STA image (i.e., image with largest CI) obtained for each cell (Fig. 4D).

**Figure 4.**
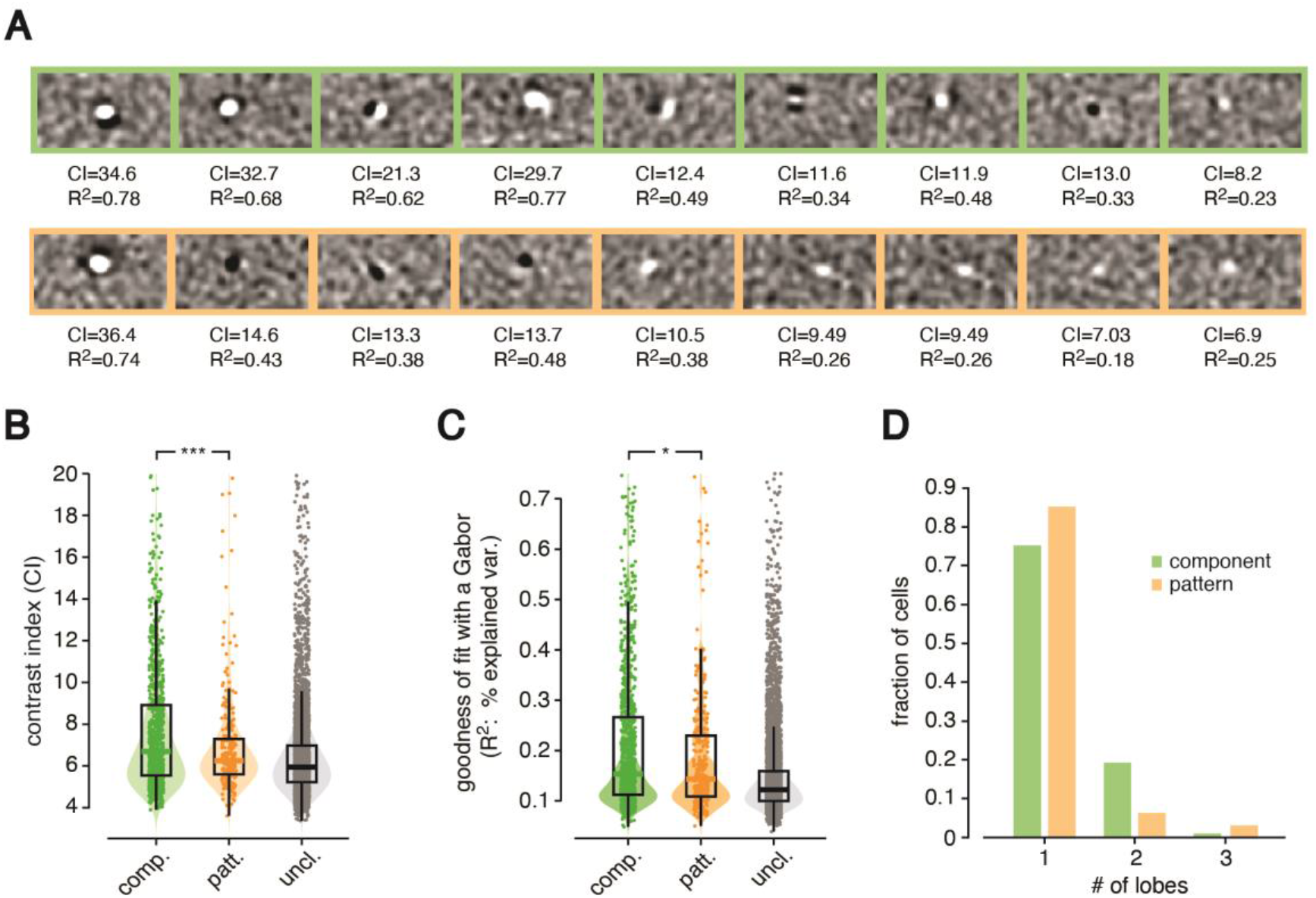
Component cells have sharper and more Gabor-like, linear receptive fields than pattern cells. (**A**) Representative examples of STA images obtained for 9 component cells (top, green frames) and 9 pattern cells (bottom, orange frames). Each image is shown along with the corresponding contrast index (CI) and a metric (R^2^) that quantifies the goodness of fit with a Gabor function. For each neuron, the RF shown here is taken at the time before spike generation when the number of distinct sub-fields (lobes) was the largest. (**B**) Distributions of CI values of the STA images obtained for the area-pooled populations of component (left, green), pattern (center, orange) and unclassified (right, gray) units at all tested (ten) time lags from spike generation. All units in the proper quadrants of the Z_c_/Z_p_ plane, with no constraint on direction selectivity, were included in this analysis (i.e., all green, orange and gray dots of Fig. 3A-C were used, no matter whether light or dark). The median CIs of component and pattern cells were significantly different according to a Wilcoxon test (\*\*\**p* = 0.003). (**C**) Same as in B, but for the distributions of R^2^ values, whose medians were significantly different, for the component and pattern cells, according to a Wilcoxon test (\**p* = 0.03). (**D**) Distribution of the number of distinct lobes in the STA image (binarized at 3.5σ) with the largest CI obtained for the component (green bars) and pattern (orange bars) cells shown in Fig. 3A-C (as in B and C, no constraint was imposed on the level of direction selectivity). The distributions were statistically different according to a χ^2^ test (*p* = 0.047).

Note that, to increase statistical power, in this analysis we counted as component and pattern cells all units in the proper quadrant of the Z_c_/Z_p_ plane, with no constraint on direction selectivity (this corresponds to all orange and green dots in Fig. 3A-C, no matter whether light or dark). This yielded a total of 98 component and 30 pattern cells, each contributing 10 STA images. As illustrated in Fig. 4B and Fig. 4C, both the median CI and GOF were slightly but significantly larger for component than for pattern cells (*p* = 0.003 and *p* = 0.03, Wilcoxon test). The lobe count distributions were also statistically different (*p* = 0.047; χ^2^ test), with a larger proportion of pattern than component cells having RFs with just a single, prominent lobe (Fig. 4D).

Overall, this indicates that STA was relatively less successful at capturing the spatial structure of RFs in the case of pattern cells. This does not mean that it completely failed at doing so. As shown by the example neurons in Fig. 2D-F and Fig. 4A, STA did often return, even in the case of pattern cells, some well-structured RFs. However, the overall lower contrast, “Gaborness” and complexity of the RFs of pattern cells suggests a more prominent contribution of nonlinear terms (not captured by STA) in establishing their stimulus-response mapping, as compared to component cells. The crucial question is the extent to which the linear RFs inferred via STA are able to account for the pattern and component nature of the two populations.

### Response predictions based on linear receptive fields yield an incorrect classification of pattern cells

For the example component cells of Fig. 2A-C, the observed tuning curves for both grating and plaid direction (solid green lines) were generally well matched by the curves predicted on the base of the linear RFs inferred via STA (dashed green lines). As a result, the units retained their classification as component cells when the Z_c_ and Z_p_ indexes were computed on the predicted curves. By contrast, for the example pattern cells of Fig. 2D-F (solid orange lines), the predictions based on linear RFs (dashed orange lines) accounted at most for the tuning for grating orientation but failed to capture the tuning for plaid direction. This led to a misclassification of two of the units as component when the Z_c_ and Z_p_ scores were computed on the predicted curves. In this section, we checked whether this phenomenon was statistically consistent at the population level.

Predictions obtained using STAs as linear filters in a LN model work at best when the spatial frequency of the stimulus fed to the model matches the dominant spatial frequency of the STA itself (i.e., the spatial scale of the excitatory and inhibitory lobes in the STA). This, in turns, depends in a nontrivial way on how the selectivity of the neuron of interest interacts with the properties of the stimulus used for reconstruction (e.g., the spatiotemporal correlated noise movies used in our study) (Sharpee et al., 2006). In our case, we empirically observed that the spatial scale of the RFs estimated via STA better matched the gratings with SF = 0.02 cycles/deg. We therefore computed predictions for responses to gratings and plaids with such spatial frequency. To guarantee the consistency with the analysis shown in Fig. 3 (where the Z_p_ and Z_c_ indexes were computed for stimuli at the most effective SF), we restricted the pool of neurons included in this analysis to units that were consistently classified as either component or pattern, regardless of the SF of the stimuli used to compute their direction tuning (e.g., units that were classified as patterns at both SF=0.04 and SF=0.02). This reduced the overall pools of component and pattern cells (merged across the three areas) from, respectively, 98 and 30 (i.e., all green and orange dots across Fig. 3A-C) to 58 and 17.

Fig. 5A illustrates the distribution of these two populations in the Z_c_/Z_p_ plane (light green and light orange dots) and shows how the pairs of Z_c_ and Z_p_ values obtained for each unit changed when it was computed on the direction tuning curves predicted by the LN model (dark green and dark orange dots). The difference between the two populations was striking. While most component cells remained in the “component” region of the plane and those that changed category became at most unclassified, none of the pattern cells retained its classification and many of them ended up being classified as component. This is better quantified by the bar plot in Fig. 5B, comparing which fraction of units in the two populations retained its original classification (e.g., component remaining component), which fraction switched to the opposite class (e.g., component becoming pattern) and which fraction became unclassified, based on their responses to gratings and plaids as predicted by the LN model. The two distributions were radically (and significantly; *p* = 5.9*10^-11^, χ^2^ test) different, thus testifying the inability of the linear receptive fields estimated via STA to capture the “patternness” of pattern cells.

**Figure 5.**
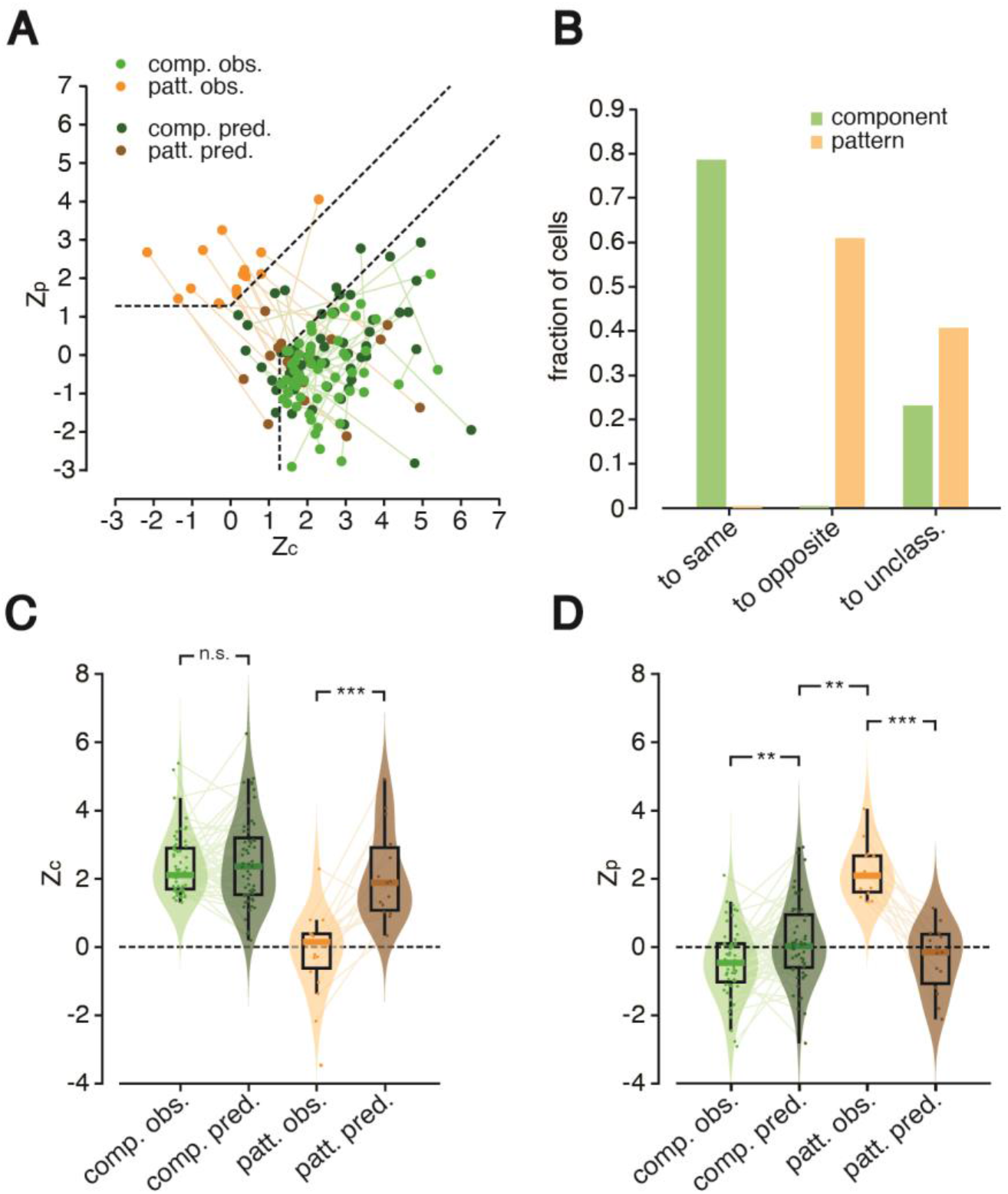
Linear receptive fields do not account for global motion selectivity of pattern cells. (**A**) Scatter plot showing the distributions of the pairs of observed Z_p_ and Z_c_ indexes for pattern (light orange) and component cells (light green) together with the values of the same indexes (dark orange and green dots), when computed on the tuning curves predicted using the spatiotemporal RFs that were estimated via STA. The lines connect the observed and predicted pairs of index values to highlight the displacement in the Z_p_/Z_c_ plane for each unit. The neuronal populations shown here include pattern and component cells pooled from all targeted areas that met the constraint to maintain the same classification (i.e., either pattern or component) at both the spatial frequencies (SF) tested in our experiment (i.e., 0.02 and 0.04 cpd). All the analyses were performed on observed and predicted responses to gratings and plaids presented at SF = 0.02 cpd. Dashed lines are decision boundaries in the Z_p_/Z_c_ plane that distinguish regions where cells are labeled as component (bottom-right corner) or pattern (top-left corner). (**B**) Bar plot reporting the fraction of units (pattern cells in orange and component cells in green) that preserved their original classification (left; “to same” label), switched to the opposite class (center; “to opposite” label) or landed in the unclassified region (right: “to unclass.” label), when the Z_p_ and Z_c_ indexes were computed on the predicted direction tuning curves. (**C**) Distributions of observed (light colors) and predicted (dark colors) Z_c_ values for the two populations of component (green) and pattern (orange) cells. (**D**) Distributions of observed (light colors) and predicted (dark colors) Z_p_ values for the two populations of component (green) and pattern (orange) cells. In C and D, the statistical comparison between the medians of the observed vs. predicted distributions for the same neuronal populations was performed using a paired Wilcoxon test (\*\**p* < 0.01; \*\*\**p* < 0.001). In D, the statistical comparison between the median of the predicted Z_p_ indexes for the component cells vs. the median of the observed Z_p_ indexes for the pattern cells was carried out using an unpaired Wilcoxon test (\*\**p* < 0.01).

To better understand the cause of this phenomenon, we separately plotted for the two cell populations (i.e., component and pattern) the distributions of the Z_c_ and Z_p_ indexes, as computed based on the observed responses (light shading) and the predicted ones (dark shading). In the case of component cells (Fig. 5C, green shadings), Z_c_ remained very stable, with no detectable difference between the medians of the observed and predicted values (*p* = 0.84, paired Wilcoxon test). By contrast, for pattern cells (Fig. 5C, orange shadings), the median Z_c_ increased substantially (and significantly; *p* < 0.001, paired Wilcoxon test): from close to zero (as it had to be, given the classification of these units as pattern cells based on their observed responses) to close to 2 (i.e., close to the value observed for component cells). This, together with the trend observed for Z_p_ (see below), explains why many pattern cells switched their status from pattern to component when the responses of the LN model were considered (see Fig. 5A-B).

A similar result was found for the Z_p_ index (Fig. 5D). In the case of component cells (green shadings), its median remained quite stable, although it was significantly closer to zero for the predicted than for the measured responses (*p* = 0.005, paired Wilcoxon test). This points to a difficulty of the LN model to fully capture, in the case of component cells, the extent to which tuning for plaid direction is anticorrelated with the one of an ideal pattern cell. Critically, however, this increase of Z_p_ was marginal and the index remained substantially and significantly lower than the large, positive values (close to 2) observed for pattern cells (compare the dark green to light orange distributions; *p* = 0.006, unpaired Wilcoxon test). This explains why most component cells retained their classification when Z_c_ and Z_p_ were computed based on their predicted responses (Fig. 5A-B). Pattern cells displayed instead a very different behavior (orange shadings). The median Z_p_ dropped to zero when computed using the predicted responses of the LN model, being substantially lower than the value (close to 2) obtained for the observed responses (*p* < 0.001, paired Wilcoxon test). This explains why none of the pattern cells retained its pattern status when the predicted responses were considered (see Fig. 5A-B).

Overall, this analysis shows that linear RFs inferred via STA are not good predictors for the global-motion sensitivity of pattern cells, when used as linear filters in a LN encoding model. As examined in the next section and in the Discussion, this result has important implications concerning the nature of pattern cells in rodent visual cortex.

### Comparison with a state-of-the-art, convolutional neural network model of dorsal stream processing

The results shown in the previous sections suggest that a fraction of rat visual cortical neurons encode global motion direction of complex visual patterns via integrative, nonlinear processes that are consistent with those thought to be at work at the higher stages of the monkey dorsal stream (Khawaja et al., 2009; Rust et al., 2006). However, a direct comparison of motion cortical processing between rats and monkeys is hard to perform, because some of the properties measured in our study (e.g., cross-orientation suppression and the ability of STA-based LN models to account for responses to plaids and gratings) have not been systematically investigated in monkeys – and vice versa, some of the processes targeted by studies of the monkey dorsal stream (e.g., tuning for the dominant motion direction of random dot kinematograms) (Born and Bradley, 2005) have not been thoroughly explored in rodents.

To bridge this gap and test the extent to which the properties of rat component and pattern cells are consistent with the existence of a functional motion processing hierarchy we carried out a comparison with DorsalNet, a state-of-the-art neural network model of the monkey dorsal stream (Mineault et al., 2021). The development of DorsalNet was inspired by the success of deep convolutional neural networks trained for image classification to model the tuning of visual neurons along the monkey ventral stream (i.e., the primate shape processing pathway) with unprecedented accuracy (Yamins et al., 2014; Yamins and DiCarlo, 2016; Zhuang et al., 2021). By applying the same rationale to the domain of motion processing, (Mineault et al., 2021) trained a 6-layer 3D convolutional neural network (named DorsalNet; see Fig. 6C) with the self-supervised learning objective of predicting the self-motion parameters of a virtual agent moving in a simulated environment from its own visual input. As a result of training, the units of the network developed a tuning for visual motion that can explain visual responses in a database of neural recordings from the primate dorsal stream better than many other computational models of motion processing. This makes DorsalNet the current, best-in-class, in-silico model of the dorsal stream.

**Figure 6.**
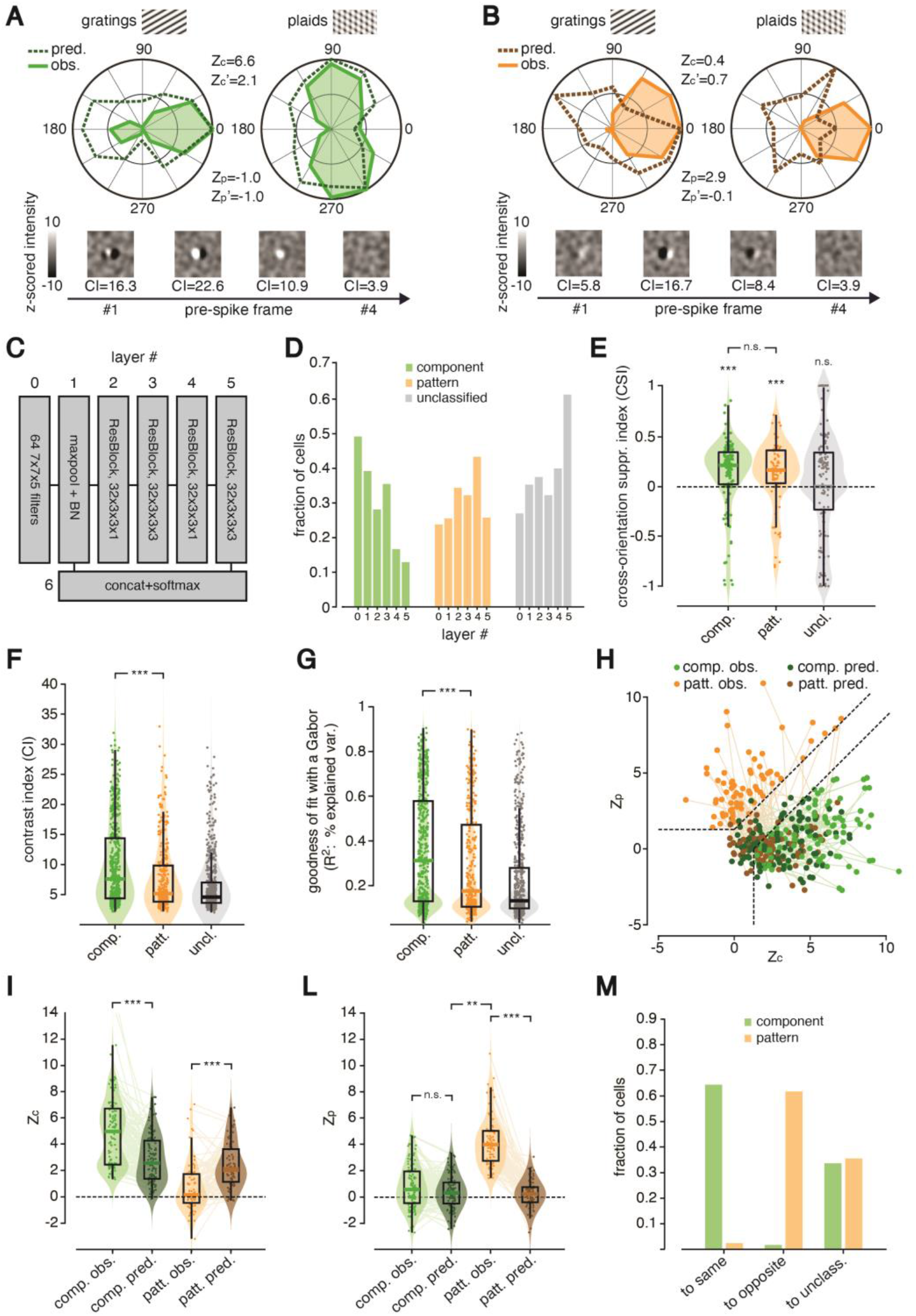
Units of DorsalNet, a neural network model of dorsal-stream processing, display tuning properties that are highly consistent with those observed for rat visual cortical neurons. (**A**) Example of a DorsalNet unit displaying the typical tuning of a component cell (same layout and color code used in Fig. 2A-C). (**B**) Example of a DorsalNet unit displaying the typical tuning of a pattern cell (same layout and color code used in Fig. 2D-F). (**C**) Schematic of DorsalNet layer structure. (**D**) Fraction of component (green bars), pattern (orange bars) and unclassified (gray bars) units from layer #0 to layer#6 of DorsalNet (compare to Fig. 3E). (**E**) Distribution of CSI values for component (green), pattern (orange) and unclassified (gray) units, pooled across all DorsalNet layers (same layout, color code and statistical analysis as in Fig. 3F). (**F-G**) Distribution of CI and R2 values (the latter assessing the goodness of fit with a Gabor function) for component (green), pattern (orange) and unclassified (gray) units, pooled across all DorsalNet layers (same layout, color code and statistical analysis as in Fig. 4B-C). (**H**) Scatter plot showing: i) the distribution of observed Z_p_ and Z_c_ indexes for pattern (light orange) and component (light green) units, pooled across all DorsalNet layers; and 2) the values of the same indexes (dark orange and green), as computed for the tuning curves predicted using the STA-spatiotemporal RFs that were estimated via STA (compare to Fig. 5A). (**I-L**) Distributions of observed (light colors) and predicted (dark colors) Z_c_ and Z_p_ values for the populations of DorsalNet units classified as component (green) and pattern (orange; same layout, color code and statistical analysis as in Fig. 5C-D). (**M**) Bar plot showing the fraction of DorsalNet units (pattern in orange and component in green) that preserved their original classification (left; “to same” label), switched to the opposite class (center; “to opposite” label) or landed in the unclassified region (right: “to unclass.” label), when the Z_p_ and Z_c_ indexes were computed on the predicted direction tuning curves (compare to Fig. 5B).

In our analysis, we used the pre-trained DorsalNet model provided by (Mineault et al., 2021), and we fed it with gratings and plaids drifting along the same 12 directions used in our recordings, as well as with spatiotemporally correlated noise movies derived from those used in our experiments. We then measured the responses (i.e., the activations) to these stimuli of all the units at the center of the convolutional map in the output layer of each of the 6 blocks of the network (Fig. 6C). Finally, we analyzed the tuning properties of the sampled units by employing the same data processing pipeline applied to the neuronal data.

As already reported by (Mineault et al., 2021), many units across the layers of the network displayed sharp direction selectivity, which, in some case, was consistent with the tuning of a component cell, while, in some other, with the tuning of a pattern cell. The example tuning curves reported in Fig. 6 A and B illustrate such cases, along with the sequences of STA images showing the inferred spatiotemporal structure of the units’ RFs (a striking similarity with the example component and pattern cells displayed in Fig. 2 can be noticed). As expected from a dorsal-like motion processing hierarchy, the proportions of component and pattern units traded off across the ResNet blocks of the network, with the former sharply decreasing from layer 0 to layer 5 and the latter smoothly increasing (compare the green to the orange bars in Fig. 6D), although both kinds of units coexisted in all layers. Interestingly, both the component and pattern units responded less strongly to plaids than to gratings showing, on average, cross-orientation suppression index values significantly greater than zero (Fig. 6E; component: *p* = 0.0001, unpaired t-test; pattern: *p* = 0.001, Wilcoxon test) but not statistically different from each other (*p* > 0.05; Wilcoxon test), as already observed for rat visual cortical neurons (Fig. 3F).

More importantly, we found that, also in DorsalNet, the spatiotemporal RFs of component units (as inferred via STA) were sharper (i.e., with higher contrast; Fig. 6F; *p* = 4.2*10-^7^ Wilcoxon test), better approximated by Gabor filters (Fig. 6G; *p* = 4.1*10^-6^ Wilcoxon test), and better capable to predict the observed Z_p_ and Z_c_ values (Fig. 6H-L; see legend for a statistical comparison of observed and predicted Z values), and, therefore, to preserve the units’ classification (Fig. 6M; *p* = 1.0*10^-300^; χ^2^ test), as compared to the RFs obtained for pattern units. All these trends matched strikingly well those observed for rat component and pattern cells (compare to Fig. 4 and 5). Only, some of them were sharper in DorsalNet, likely due to the noiseless nature of the activations of the units of the network, as compared to neuronal responses. In particular, the consistency between the properties of pattern cells in rat visual cortex and those of pattern units in DorsalNet strongly supports the conclusion that the former are the result of non-trivial, hierarchical computations aimed at integrating local motion signals into pattern-invariant representations of global motion direction.

## Discussion

The first goal of our study was to measure the incidence of pattern and component cells in some visual cortical areas of the rat that, by analogy with previous mouse studies (Juavinett and Callaway, 2015; Marshel et al., 2011; Muir et al., 2015; Palagina et al., 2017), are expected to be involved in dorsal stream processing: V1, LM and RL. Our results (Fig. 3A-C) are in agreement with two mouse studies showing a sizable fraction of neurons that are sensitive to global motion direction in V1 (Muir et al., 2015; Palagina et al., 2017), while they are at odd with another study reporting no pattern cells in this area (Juavinett and Callaway, 2015). At the same time, they support the conclusion of the latter study that mouse LM is rich in pattern cells, but are in disagreement with the finding of an abundance of pattern cells in RL (Juavinett and Callaway, 2015).

One possible explanation for such discrepancies is the different kind and dosage of anesthetic used during the recordings. In fact, it is well established that pattern cells are almost absent from V1 of anesthetized monkeys (Khawaja et al., 2013, 2009; Movshon et al., 1985; Movshon and Newsome, 1996), while (Guo et al., 2004) reported a sizable amount of this cell class (9%) in awake macaques. Such responses were likely enabled by feedback from higher level areas that are cut off during anesthesia. This suggests that, in principle, the level and kind of anesthesia can affect motion integration computations in V1. Consistently, (Palagina et al., 2017), who did observe pattern cells in V1, used lower isoflurane concentrations (0.6%) than (Juavinett and Callaway, 2015) (from 0.6 to 1.2%), who did not. Interestingly, (Muir et al., 2015), who used a fentanyl and medetomidin anesthesia similar to the one employed in our study, reported proportions of pattern and component cells in V1 (respectively, 3% and 31%) that are very close to those found in our experiments (6% and 27%; Fig. 3E).

While these considerations could reconcile the contrasting results concerning the presence of pattern cells in V1 of rodents, the disagreement between (Juavinett and Callaway, 2015) and our study on the incidence of pattern cells in RL could underlie possible differences on the functional role of this area between rats and mice. At the same time, it is worth pointing out that our results are consistent with a recent mouse study, where wide-field calcium imaging was used to the investigate responses to coherent motion stimuli across the dorsal cortex of head-fixed mice (Sit and Goard, 2020). The authors found that, on average, responses to coherent motion stimuli (i.e., RDKs) are strong in the extrastriate areas that are located medially to V1 (i.e., AM and PM), as well as, laterally to V1, in AL and in the most anterior and lateral part of LM. On the other hand, they reported a clear lack of strong coherent motion responses in RL, which suggests that this area may not be specialized for the processing of pure visual motion or optic flow information (differently, e.g., from monkey MT). Taken together with recent evidence showing that RL neurons are particularly attuned to near field visual stimuli (La Chioma et al., 2019), as well as with the privileged position of this area, as a part of posterior parietal cortex (PPC) (Lyamzin and Benucci, 2019), to act as a hub of visuotactile integration and multimodal decision making (Licata et al., 2017; Nikbakht et al., 2018; Olcese et al., 2013; Raposo et al., 2014), our results support the hypothesis that RL plays a functional role that is different from pure, high-level processing of visual motion information. On the other hand, the existence of pattern cells in such a low-level area as V1 corroborates the notion that, in rodents, primary visual cortex might contribute to motion-related computations that, in monkeys, are carried out by higher-level areas like MT (possibly reflecting the shallower hierarchy of the visual system of mice and rats). Interestingly, this is consistent with a recent study (Tohmi et al., 2021) reporting the presence of V1 neurons that are selective for motion-streak (i.e., the smeared representations of fast-moving stimuli arising from temporal integration) – another, mid-level computation of motion information that, in monkeys, is performed by MT neurons (Albright, 1984).

The second goal of our study was to assess the extent to which cross-orientation suppression affects component and pattern cells in rat visual cortex. Addressing this question is important for two reasons. First, cross-orientation suppression belongs to a class of nonlinear tuning phenomena that are thought to depend on divisive normalization, a canonical cortical computation where the response of a neuron is divided (normalized) by the summed activity of a pool of nearby neurons. Divisive normalization has been called into cause to explain a variety of nonlinear interactions among competing stimuli within the classical and extraclassical receptive fields of visual neurons in cats and monkeys (Carandini and Heeger, 2012). There is now a strong interest in establishing whether the same interactions take place in rodent visual cortex, because of the potential of dissecting the underlying neuronal circuitry using the molecular tools that rodent studies allow (Niell and Scanziani, 2021). Several studies have succeeded at demonstrating the existence of contrast tuning and surround modulation in mice, examining its dependence from different classes of inhibitory neurons (Adesnik, 2017; Adesnik et al., 2012; Ayaz et al., 2013; Keller et al., 2020; Niell and Scanziani, 2021; Samonds et al., 2017; Self et al., 2014; Vaiceliunaite et al., 2013). Cross-orientation suppression is comparatively less explored in rodents (but see (Alwis et al., 2016)). Our study is one of the first to show that this normalization process equally affects the responses of rat visual cortical neurons, regardless of whether they are component or pattern cells, deepening our understanding this phenomenon in the context of visual motion processing.

This result is also important because it allows ruling out one of the mechanisms we have previously hypothesized to account for the observed asymmetry in the ability of rats to generalize a motion discrimination task from grating to plaids or vice versa. As explained in (Matteucci et al., 2021), if cross-orientation suppression affected more strongly component than pattern cells, downstream decision neurons would learn to rely mainly on the latter to represent motion direction, in case of training with plaids. This would allow a good generalization when the animals are tested with gratings (given the consistency of the tuning of pattern cells with the two stimulus classes). In case of training with gratings, decision neurons would instead learn to rely on both component and pattern cells, but, being the former way more numerous (as also shown in this study), the resulting representation would not guarantee a good generalization from gratings to plaids. Our recordings, by showing that both cell types are equally affected by cross-orientation suppression, rules out this scenario and suggests instead an alternative scheme where component cells act as the building blocks that are necessary to “assemble” pattern cells, but only the latter are read out by decisions neurons to infer motion direction. As explained in (Matteucci et al., 2021), this is a scenario where pattern cells sit higher than component cells along the rat functional motion processing hierarchy, consistently with the role that area MT and MST play with respect to area V1 in monkeys (Khawaja et al., 2009). Under this hypothesis, a good generalization can be achieved from training with plaids to testing with gratings but not vice versa, because of the different magnitude of the synaptic inputs from pattern cells to decision neurons that would result from training the animal with gratings or plaids (see Fig. 6 in (Matteucci et al., 2021)). Our recordings are consistent with this scenario, thus suggesting that rodent pattern cells, despite their scant number and their incidence in multiple visual areas, plays a similar functional role to their homologues in monkey MT. This conclusion is consistent with the well-established ability of rodent visual cortical neurons to send (receive) functionally specific inputs to (from) downstream (upstream) areas (D’Souza et al., 2019; Gao et al., 2010; Glickfeld et al., 2013; Ji et al., 2015; Matsui and Ohki, 2013). Interestingly, a recent study performing retrograde and anterograde tracing as well as optogenetic-based connectivity assessment evidenced a clear preferential connectivity between LM and anterior cingulate cortex (ACC) (Sidorov et al., 2020) – an area that has been recently proposed as a key node for visuomotor transformation in mice (Kim et al., 2021). These results, taken together with our finding that LM is an area rich of pattern cells (Fig. 3E), strengthen the preferential connectivity hypothesis explained above. Projection-specific or axonal imaging experiments targeted at functionally characterizing the input from V1 and LM to higher-order areas involved in perceptual decision making such as ACC, M2 (Yang and Kwan, 2021) or PPC (Licata et al., 2017; Nikbakht et al., 2018; Odoemene et al., 2018;) could test directly this hypothesis, by verifying whether the populations of projecting neuron relaying visual information to this areas are particularly enriched in pattern cells.

The last and more important goal of our study was to assess whether pattern cells in rodents perform a truly integrative, nonlinear processing of local motion signals to encode global motion direction of complex patterns, as the plaid stimuli. Answering this question is important because, as originally pointed out by (Tinsley et al., 2003) and shown by the example we provided in Fig. 1, tuning consistent with pattern-like behavior may emerge from “blobby”, purely linear, Gabor-like RFs with low aspect ratio. Following a similar argument, also (Palagina et al., 2017), having observed that the classification of the recorded V1 units into the pattern and component classes was sensitive to the aperture angle of the plaid stimuli, suggested that pattern responses in mouse V1 could rely more on the geometry of the receptive field (i.e. matching of local contrasts with RF’s on/off-subfields; see Fig. 1C) rather than the nonlinear process thought to be at work in primates (Rust et al., 2006).

To address this issue, we applied reverse correlation analysis (STA) to reconstruct the linear RF of units classified as either component or pattern cells by processing their responses to spatiotemporally correlated noise movies (see Materials and methods for details). As expected, for many component cells, we obtained temporal sequences of high-contrast STA images (Fig. 4A-B), typically Gabor-like (Fig. 4C), often containing a pair of flanking lobes with opposite polarity (i.e., one excitatory and the other inhibitory; Fig. 2, 4A and 4D). In general, these STA images, when used as the linear kernels in a NL model, predicted the tuning of component cells for gratings and plaids well enough to keep the resulting Z_p_ and Z_c_ index values close to those observed empirically (Fig. 5A). As a result, most of component cells retained their classification when the Z_p_ and Z_c_ indexes were computed on the responses predicted by the model (Fig. 5B). Interestingly, also pattern cells yielded in many cases well-structured STA images, with a quality that, in general, was only slightly lower than that of the STA images obtained for component cells (Fig. 4). However, the ability of STA-based NL models to account for direction tuning was strikingly poorer for pattern cells – none of the neurons classified as pattern on the ground of their observed responses retained its classification when the Z_p_ and Z_c_ indexes were computed on the responses predicted by the model (Fig. 5A-B). This suggests that, although STA can capture some linear “residue” of the spatiotemporal selectivity of pattern cells, it is not able to account for their tuning. This rules out the possibility that the pattern responses we observed (e.g., see Fig. 2D-F) are the result of linear filtering processes performed by “blobby” RFs (of the kind illustrated in Fig. 1C). On the contrary, our results suggest that the computations carried out by pattern cells to encode motion direction rely on nonlinear processes that STA (by construction, being a linear method) cannot capture.

This conclusion has important implications for our understanding of the nature of motion processing in rodent visual cortex. Looking at the primate dorsal stream literature, the most established mechanistic models of pattern-motion selectivity in MT are based on the nonlinear pooling of inputs from narrowly tuned, V1-like component cells, whose preferred motion directions are spread over a wide range of angles (Rust et al., 2006). We still do not know the extent to which such models can be extended to rodents. Certainly, they will need to be adapted to fit the specificities of the rodent visual system – e.g., to account for the larger contribution of direction selective inputs from retinal ganglion cells (Rasmussen et al., 2020) and the overall broader tuning of LGN and intracortical inputs (Durand et al., 2016; Gao et al., 2010; Matteucci et al., 2019; Niell and Stryker, 2008; Palagina et al., 2017). Our study, however, suggests that also in rodents the tuning of pattern cells is mainly determined by truly nonlinear, integrative processes, possibly homologous to the ones at work in primates and instantiated in the above-mentioned models.

To test further this hypothesis, it would be interesting to compare our findings to similar analyses performed on monkey pattern cells. Unfortunately, we are not aware of monkey studies in which STA was applied to characterize the structure of component and pattern cells and then to predict their direction tuning, as done here. To our knowledge, the only reverse correlation study in which the RF structure of MT neurons was mapped using very sparse noise did not differentiate between component and pattern cells, and neither measured nor predicted neuronal responses to gratings and plaids (Livingstone et al., 2001). Similarly, although several monkey studies report examples of tuning curves of component and pattern cells for grating and plaid direction (Rust et al., 2006; Smith et al., 2005), we are not aware of a systematic comparison of the impact of cross-orientation suppression on the two cell classes in the monkey literature.

This is one of the reasons we performed a comparison with a state-of-the-art, neural network model of the dorsal stream: DorsalNet (Mineault et al., 2021). Our goal was to compare the tuning properties of component and pattern cells recorded from rat visual cortical areas to those of the units of a benchmark computational architecture, where selectivity for increasingly complex motion patterns is built via hierarchical processing. As shown by (Mineault et al., 2021), the selectivity of neurons sampled from progressively higher stages of the monkey dorsal stream is better explained by the activations of units located at progressively deeper layers of DorsalNet – i.e., layers 1, 2, and 3 best match areas V1, MT and MST. Our analysis of the tuning of DorsalNet units for gratings and plaids corroborated this conclusion, showing that, while the proportion of component units decreases along the hierarchy, the fraction of pattern units increases (Fig. 6D) – the same trend found along the monkey dorsal stream (Khawaja et al., 2009). Together, these findings indicate that DorsalNet successfully captures some of the core hierarchical processes underlying motion representation along the monkey dorsal stream. This makes DorsalNet extremely valuable as a benchmark against which to compare the properties of a system, such as rodent visual cortex, whose level of sophistication in terms of motion processing is still poorly understood, at least as compared to primate visual cortex.

Our analyses show that three key phenomena we observed in the rat visual cortex also occur in DorsalNet: 1) the presence of both component and pattern units, with an increase in abundance of the latter in higher stages of processing, possibly reflecting what we observed between V1 and LM (compare Fig. 6D to Fig. 3E); 2) the similar impact of cross-orientation suppression on these two classes of units (compare Fig. 6E to Fig. 3F); and 3) the success of linear spatiotemporal filters inferred via STA to account for the tuning (and, therefore, the classification) of component units but not of pattern units (compare Fig. 6H-M to Fig. 5). The latter result is particularly important. In fact, finding the same evidence for integrative, nonlinear computations being performed by rat pattern cells and DorsalNet pattern units further corroborates a potential homology with the monkey dorsal stream.

In summary, our experimental findings, supported by the comparison with the tuning properties of DorsalNet units, provide compelling evidence for the existence of truly advanced encoders of global motion direction distributed across rat visual cortical areas V1 and LM. A wide range of molecular and genetic tools are available in rodents to study the detailed mechanisms underlying sensory coding properties such as direction selectivity. These tools have already been used, for instance, to successfully unravel the neural circuit motif underlying direction selectivity in mouse V1 (Rossi et al., 2020). Our findings set the stage to exploit these tools to dissect the circuit-level mechanisms underlying integration of local motion signals into the representation of global motion direction. At the same time, our study provides yet another example of the important insights that deep neural network models can yield about the processing performed by the rodent visual system and the extent to which such processing is comparable to that carried out by the monkey ventral and dorsal streams (Matteucci et al., 2019; Muratore et al., 2022).

## Materials and methods

### Surgery and recordings

All animal procedures were in agreement with international and institutional standards for the care and use of animals in research and were approved by the Institutional Animal Care and Use Committee of the International School for Advanced Studies (SISSA) and by the Italian Ministry of Health (project DGSAF 22791-A, submitted on 7 September 2015 and approved on 10 December 2015, approval 1254/2015-PR).

We performed extracellular neuronal recordings from 29 naïve male Long Evans rats, weighted 300-700 grams and aged 3-12 months. Each rat was anesthetized with an intraperitoneal (IP) injection of a solution of 0.3 mg/kg of fentanyl (Fentanest®, Pfizer) and 0.3 mg/kg of medetomidin (Domitor®, Orion Pharma). During the surgery, we monitored the anesthesia level by checking the animal paw reflex and by measuring the oxygenation and heart rate through a pulse oximeter (Pulsesense-VET, Nonin). To avoid anesthesia-induced hypothermia, temperature was monitored and maintained at 37° C through a heating pad. To prevent hypoxia, a constant flux of oxygen was delivered to the animal throughout the surgery. A constant level of anesthesia was maintained by continuously delivering an IP injection of the same aesthetic solution used for the induction, but at a lower concentration (0.1 mg/kg/h fentanyl and 0.1 g/kg/h medetomidin), by means of a syringe pump (NE-500; New Era Pump Systems). Once deeply anesthetized, the animal was secured to a stereotaxic apparatus (Narishige, SR-5R) and we performed a craniotomy on the left hemisphere, over the selected target (typically, a ~ 4 mm^2^ window). Stereotaxic coordinates of the center of the craniotomy were 6.5 mm posterior from bregma and 4 mm left to the sagittal suture (i.e., anteroposterior or AP, 6.5; mediolateral or ML, 4.0) for sessions targeting V1, 7 mm AP and 5 mm ML for sessions targeting ML and 5 mm AP and 4.5 mm ML for sessions targeting RL.

To keep eyes hydrated during the surgery, we protected them by applying an ophthalmic ointment (Epigel®, Ceva Vet). Once completed the surgery, the rat was placed over a rotating platform, with the right eye just in front of the center of the screen (distance = 30 cm) and the left eye covered with non-transparent black tape. The right eye was fixed through a metal eye-ring to prevent eye movements during the visual stimulation protocol.

Extracellular recordings were performed under light anesthesia while rats were passively exposed to visual stimulation. Recordings were carried out using either single- (or double-) shank 32- (or 64-) channel silicon probes (NeuroNexus Technologies) with a site recording area of 775 μm^2^ and an inter site spacing of 25 μm. The insertion of the electrode into the cortex was performed through an oil hydraulic micromanipulator (Narishige, MO-10). The insertion depth was different for each area: for V1 and RL, it was ~900 μm with an insertion angle relative to the cortical surface of ~20°; for LM, it was ~1500 μm, with an insertion angle of ~25°. Neuronal signals were recorded with a sampling rate of 25 kHz, and were acquired and pre-amplified using a system three TDT (Tucker-Davis Technologies) workstation.

### Single-unit isolation

Single-units were isolated offline using the KlustaKwik-Phy software package (Rossant et al., 2016). After the automatic spike detection and features extraction, we performed a manual refinement of the sorting through the “Kwik-GUI” interface. The manual refinement of the automatic output was sorted based on the following criteria: i) the compactness of the clusters in the space of the principal components of the waveforms; ii) the shape of the auto- and cross-correlogram (the latter was used to decide whether to merge or not two clusters); iii) the variation of the principal components of the waveform over time – this was especially useful to take into account possible electrode drifts; iv) the shape of the average waveform – this was especially useful to detect artefacts or non-physiological signals. When we suspected that a cluster included spikes fired by multiple units, we ran the “reclustering” function of the GUI, in an attempt to split it into its component single units. To be included in the analyses presented in the Results, single units were required to meet the following criteria: i) show a clear refractory period (i.e., less than 0.5% of the spikes present in <2 ms of the spikes’ autocorrelogram); and ii) be clearly grating or plaid responsive, i.e., with the response to the most effective grating or plaid condition being larger than 2 spikes/s (baseline-subtracted) and being larger than six z-scored points relative to baseline activity. The average baseline (spontaneous) firing rate of each well-isolated unit was computed by averaging its spiking activity over every inter stimulus interval.

### Visual stimuli

During each recording session, two kinds of visual stimulation protocols were administered to the rats. The first one was a RF mapping procedure, aimed at estimating the average preferred retinotopic location of the units recorded at each site along the length of the probe. This procedure allowed identifying the visual area each unit was recorded from, by tracking the reversals of the retinotopy that, in rodents, take place at the boundaries between adjacent visual areas. The details of this procedure have been previously described in several other studies from our group (Matteucci et al., 2019; Piasini et al., 2021; Tafazoli et al., 2017). This RF mapping protocol consisted in the presentation of 10° long drifting bars spanning different orientations (0°, 45°, 90°, 135°). Each bar, a white rectangle over a black background, was presented in 66 different positions, across a grid of 6 rows (spanning vertically 50°) and 11 columns (spanning horizontally of 100°). During the recording, we plotted in real-time the multi-unit-activity (MUA) collected at each site of the probe, so as to identify the reversal of the retinotopy (i.e., RF centers “moving” from nasal to temporal instead than from temporal to nasal) that takes place in the transition from V1 to LM or RL (Espinoza and Thomas, 1983; Marshel et al., 2011; Montero, 1993; Wang and Burkhalter, 2007). The total duration of the mapping protocol was about 15 minutes.

The second visual stimulation protocol included all the stimuli used to characterize neurons as pattern or component cells and to reconstruct their linear RFs, as described in the main text of this paper. The protocol consisted of (i) 20 repetitions (trials) of 1.5-s-long, full-contrast drifting gratings, made of all possible combinations of two spatial frequencies (0.02 and 0.04 cycles per degrees, or cpd), two temporal frequencies (2 and 6 Hz), and 12 directions (from 0° to 330°, in 30° increments); (ii) 20 repetitions of 1.5-s-long drifting plaids (made of two superimposed, half-contrast drifting gratings with a 120° cross-angle), again spanning all possible combinations of two spatial frequencies (0.02 and 0.04 cycles per degrees, or cpd), two temporal frequencies (2 and 6 Hz), and 12 directions (from 0° to 330°, in 30° increments); and (iii) 20 different 60-s-long spatially and temporally correlated, contrast modulated, noise movies (Matteucci et al., 2019; Matteucci and Zoccolan, 2020; Niell and Stryker, 2008). All stimuli were randomly interleaved, with a 1-s-long inter stimulus interval, during which the display was set to a uniform, middle-gray luminance level. To generate the movies, we started from sequences of independent white noise frames and we introduced spatial correlations by convolving them with a Gaussian kernel having full width at half maximum (FWHM) of 12.5°. Temporal correlation was introduced by convolving the movies with a causal exponential kernel with a 33-ms decay time constant. To prevent adaptation, each movie was also contrast modulated using a rectified sine wave with a 10-s period from full contrast to full contrast (Matteucci and Zoccolan, 2020; Niell and Stryker, 2008). Stimuli were generated and controlled in MATLAB (MathWorks) using the Psychophysics Toolbox package and displayed with gamma correction on a 47-inch LCD monitor (SHARP PNE471R) with 1920 × 1080– pixel resolution, a maximum brightness of 220 cd/m2, and spanning a visual angle of 110° azimuth and 60° elevation (placed at 30 cm from the eye of the animal). Grating stimuli were presented at 60-Hz refresh rate, whereas noise movies were played at 30 Hz.

### Classification of pattern and component responses

To classify a neuron as “pattern” or “component”, as done in Fig. 3, the first step was to select direction selective units among the recorded neurons. To this end, we quantified direction selectivity of single units in each area by computing the Direction Selectivity Index (DSI):

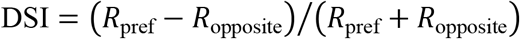

where *R*_pref_ is the response of the neuron to the preferred direction and *R*_opposite_ is the response to the opposite direction. Note that for all the analyses described here, the response of a neuron to a stimulus in a given direction x (i.e., *R*_x_) is defined as the trial-averaged, firing rate computed over entire stimulus presentation window and z-scored with respect to the spontaneous activity, as computed during all interstimulus intervals. Neurons with a DSI > 0.33 were categorized as direction selective. For the analysis aimed at comparing the linear receptive field structure of pattern and component cells using the STA method (Fig. 4) and obtain a prediction of their tuning curves using the STA images as linear kernels in a LN model, the requirement of direction selectivity was relaxed, so as to reach greater statistical power when dealing with such a rare cell type as pattern cells.

Classification of neurons as “pattern”, “component” or “unclassified” was based on their z-scored, Fisher-transformed, partial correlation indexes (Z_p_ and Z_c_), as defined as in (Movshon et al., 1985; Scannell et al., 1996):

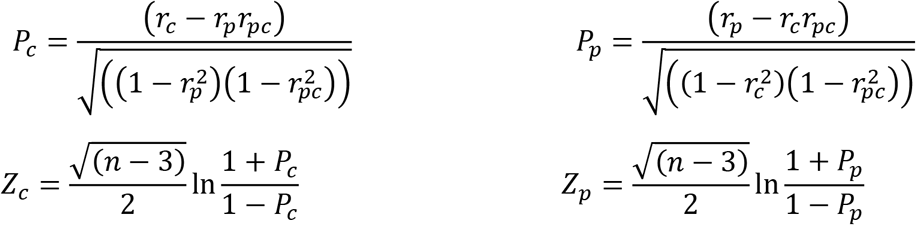

Here: i) *r_c_* is the correlation coefficient between the observed tuning curve for the plaids and the predicted tuning curve under the assumption that the unit behaves as an ideal component cell; ii) *r_p_* is the correlation coefficient between the observed tuning curve for the plaids and the predicted tuning curve under the assumption that the unit behaves as an ideal pattern cell; iii) *r_pc_* is the correlation coefficient between the two predicted curves; and iv) *n* is the number of elements of the tuning curves. The ideal pattern-cell behavior is simulated by imposing that the tuning curve is the same in response to both gratings and plaids – i.e., the predicted tuning curve for the plaids is trivially obtained by setting it identical to the tuning curve obtained for the gratings. The ideal component-cell behavior is simulated by imposing that the unit, when presented with a plaid, responds to its constituent gratings in the same way as it would respond if the gratings were presented in isolation – i.e., the predicted tuning curve for the plaids is obtained by shifting both leftwards and rightwards of half plaid cross-angle the tuning curve measured for the gratings and then averaging the two shifted versions.

At 90% confidence (as usually done in the literature) the Z critical value of 1.28 defines the pattern and component regions of the (Z_c_, Z_p_) plane, as follows:

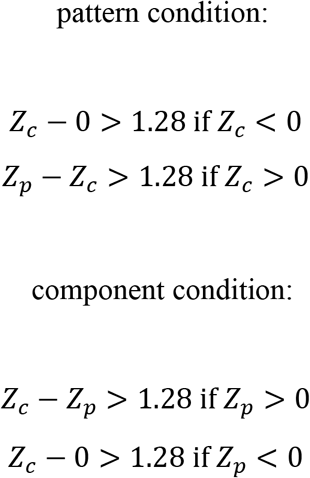

Neurons falling in these regions are classified, respectively, as pattern or components. Neurons that do not meet these criteria are considered unclassified.

### Linear RF reconstruction and prediction of pattern and component responses

Reconstruction of linear RFs underlying the selectivity of the recorded neurons was achieved employing the Spike-Triggered Average (STA) technique (Aljadeff et al., 2016; Schwartz et al., 2006; Sharpee, 2013). The method was applied to the spike trains fired by the neuron in response to the spatiotemporally correlated noise movies (see above). The STA method yields an ordered sequence of images (i.e., spatial filters), each representing the average of the stimulus ensemble at a given time lag from spike generation. STA images can therefore be interpreted as the (linear) spatio-temporal RF of a given neuron.

As we did in previous studies (Matteucci et al., 2019; Matteucci and Zoccolan, 2020), we took into account the correlation structure of our stimulus ensemble (determined by the spatiotemporal correlations in the noise movies) to reduce artefactual blurring of the reconstructed filters that such correlation could induce. To this end, we decorrelated the STA images by dividing them by the covariance matrix of the whole stimulus ensemble (Schwartz et al., 2006), using Tikhonov regularization to handle covariance matrix inversion.

Statistical significance of STA images was assessed pixelwise, by applying the following permutation test. After randomly reshuffling the spike times, the STA analysis was repeated multiple times (*n* = 30) to derive, for each pixel, a distribution of intensity values under the null hypothesis of no linear stimulus-spike relationship (i.e., random spikes being fired in a fully uncorrelated way with respect to the visual input). This allowed z-scoring the actual STA intensity values using the mean and standard deviation of these null distributions independently for each pixel. The temporal span of the spatiotemporal linear kernel reconstructed via STA extended from the time of spike generation up to 330 ms earlier. This corresponds to a duration of 10 frames of the noise movie played at 30 Hz. These procedures were performed on downsampled noise frames (16×32 pixels). The resulting STA images were then spline interpolated at higher resolution for ease of visualization and analysis, as well as to derive the subsequent LN predictions (see next paragraph).

To derive linear predictions for the grating and plaids tuning curves, as expected given the reconstructed STA image sequences, we used the latter as input stage filters of a classical linear-nonlinear (LN) model (Schwartz et al., 2006). Specifically, to obtain a prediction of the response to a stimulus drifting in a given direction, the sequence of frames of the stimulus movie was fed as an input to the STA-estimated linear filter. The output of the filter was then passed through a rectified linear unit (a.k.a. relu) to obtain the final response of the model to each stimulus frame. In the analysis presented in Fig. 5, this was done for the stimuli with SF = 0.02 cpd and taking the preferred TF for each neuron (i.e., 2 or 6 Hz). This choice was dictated by the need of carrying out the analysis with gratings and plaids matching at best the dominant spatial frequency of the reconstructed linear receptive fields, so as to maximize the quality of predictions. For this reason, the analysis reported in Fig. 5 was carried out on the subset of pattern and component neurons maintaining at SF = 0.02 cpd their pattern or component classification (determined, in the analysis of Fig. 3, at their preferred SF, i.e., either 0.02 or 0.04 cpd). To allow overfitting and implement a test/train split, the gain of the nonlinear activation function was determined, for each neuron, by fitting the LN model to the responses of the unit to the gratings presented at the non-preferred TF (i.e., the fit was not carried out on the responses to the stimulus conditions analyzed in Fig. 5). The fit was performed using the MATLAB “fminunc” function. When carrying out the analysis on DorsalNet units, the gain was instead fixed to one (being this the gain of the relu functions implemented in network). Once obtained a predicted response to the drifting stimulus under consideration, this was averaged in time over the stimulus presentation window to get the final predicted response rate of the unit for that stimulus. Repeating this process for each direction of a given stimulus class (i.e., gratings of plaids) yielded predicted tuning curves, as the one shown in Fig. 2 and Fig. 6A-B (dashed lines). These curves were then used to re-compute the Z_p_ and Z_c_ indexes (see previous section) and (re)classify each unit as “component”, “pattern” or “unclassified” (see Fig. 5 and Fig. 6H-M).

### Linear RF structure quantification

To estimate the amount of signal contained in a given STA image, we used the “contrast index” (CI) metric that we have introduced and applied in previous studies (Matteucci et al., 2019; Matteucci and Zoccolan, 2020). Briefly, the CI is designed to be a robust measure of maximal local contrast in any STA image. Having expressed the intensity values of each STA image in terms of z-scores (against the spike-time shuffled null distributions; see previous section), the CI value is defined as the maximal local peak-to-peak (i.e., white-to-black) difference in the STA, expressed in sigma units. The local peak-to-peak distance computation was performed using MATLAB “rangefilt” function. The locality of the CI is determined by the size of the neighborhood used by rangefilt. In our analyses, this parameter was set to be 20% of the lower dimension (the height) of the reconstructed STA filter. For the analysis reported in Fig. 4, all STA images for each neuron in the component, pattern, unclassified categories were included. As in done our previous studies (Matteucci et al., 2019; Matteucci and Zoccolan, 2020), we also characterized the structural complexity of the linear RFs yielded by STA by counting the number of excitatory/inhibitory lobes that were present in a STA image. The procedure is similar to the one described in (Matteucci et al., 2019). Briefly, we applied a binarization threshold over the modulus of the z-score values of the image (at 3.5 units of standard deviation). We then computed the centroids of the simply connected regions within the resulting binarized image (i.e., the candidate lobes). Last, we applied a refinement procedure, which is detailed in (Matteucci et al., 2019), to prune spurious candidate lobes (i.e., lobes smaller than 5% of the width of the reconstructed STA filter were rejected). As a final quantification of the linear RF structure of the recorded units, we fitted a Gabor function to each STA image (using the “fit2dGabor” function by Gerrit Ecke: https://www.mathworks.com/matlabcentral/fileexchange/60700-fit2dgabor-data-options) and we evaluated the goodness of this fit by computing the coefficient of determination R^2^ (as reported in Fig. 4C and 6G).

### Quantification of cross-orientation suppression

To quantify the amount of suppression (or enhancement) of the response of each unit to the plaid as compared to the grating stimuli, we defined a normalized cross-orientation suppression index (CSI) as:

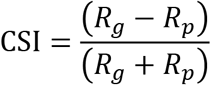

Here, *R_p_* and *R_c_* indicate the peak responses to plaids and gratings respectively (i.e., the responses to the two stimuli when presented at the most effective directions). This index takes a value of 1 for a unit responsive to gratings but not to plaids (i.e., extreme suppression) and, vice versa, a value of −1 for a unit responsive to plaids but not to gratings (i.e., extreme enhancement), whereas it takes a value of 0 for a unit showing the same peak response firing rate for both gratings and plaids. To correctly interpret this index, attention should be paid to the fact that both gratings and plaids were presented at full contrast. This was done to elicit stronger responses and increase the yield of the recordings, as well as to match the contrast of the stimuli used in our previous behavioral study (Matteucci et al., 2021). A more rigorous assessment of cross-orientation suppression would have required matching the contrast of the constituent gratings of the plaids to the contrast of the gratings presented in isolation (i.e., ideally, isolated gratings should have been presented at half contrast and the plaids at full contrast). However, since the purpose of our study was to compare the level of suppression between the two categories of component and pattern cells, and since both cell classes were probed with the same grating and plaid stimuli, our conclusions are not affected by the fact that the latter were both displayed at the same, maximal contrast. More importantly, the main goal of measuring the level of cross-orientation suppression in the two cell classes was to test which of the two mechanisms of motion integration we proposed in (Matteucci et al., 2021) can better account for rat perception of plaid motion direction – hence, the need of matching the contrast of the stimuli used in the two studies.

### DorsalNet simulations

To help interpreting the results of our analyses of visual neuronal responses, we decided to compare them with the those obtained by applying the same analysis pipeline to a state-of-the-art computational model of dorsal processing. To this aim, we choose DorsalNet, a 6-layer 3D convolutional neural network recently proposed as the best-in-class in-silico model of the dorsal stream (Mineault et al., 2021). DorsalNet is a 3D ResNet (He et al., 2016) trained with the self-supervised learning objective of predicting the parameters of simulated self-motion of an agent moving in a simulated environment from its own visual input. The training dataset consisted of a set of short videos (10 frames) of self-motion generated with the AirSim package (Shah et al., 2018), a drone and land vehicle simulation software in Unreal Engine. These videos contained simulated walking along linear trajectories with constant head rotations in two environments with starts at random positions, varying environmental conditions, lighting, starting head poses and walking speed (see (Mineault et al., 2021) for further details). Mineault and colleagues demonstrated that DorsalNet units’ activations in response to different visual stimuli can explain visual responses in a database of neural recordings along the primate dorsal stream better than many other computational models of motion processing.

In our study, we used the pre-trained DorsalNet model checkpoint provided by (Mineault et al., 2021) for our simulated experiments (“airsim_dorsalnet_batch2_model.ckpt-3174400-2021-02-12 02-03-29.666899.pt”). We fed to the network drifting gratings and plaids moving in 12 equispaced directions at a fixed spatial and temporal frequency (0.0625 pixel^-1^ and 0.0625 frames^-1^ respectively). This allowed building direction tuning curves for grating and plaid responses and classify the units of the network as pattern or component, using the same criteria applied to neuronal recordings. We also fed spatiotemporally correlated noise movie, similar to those used in our neurophysiology experiments (again, spatial and temporal correlation was achieved by filtering withe noise frames with Gaussian filters and an exponential kernel). The spatial correlation scale (i.e., sigma of the Gaussian kernel) was chosen to have a FWHM corresponding to half of the above-mentioned gratings period. This was done to ensure a good match between the spatial scale of gratings and plaids and the one of the filters inferred via STA. The time constant of the temporal kernel was chosen to have a FWHM of 1.5 frames, so as to boost the effectiveness of the noise movies in activating the units, without inducing strong temporal correlations in the inferred STA filters. Activations were sampled from all units at the center of the convolutional maps of all feature maps at the output layer of each block of the network (named layer # 1,2,3 & etc. in the main text), including the non-ResNet blocks (see Fig. 6C). This was done using the functions provided by (Mineault et al., 2021) along with the pre-trained model.

## Acknowledgments

This work was supported by a European Research Council Consolidator Grant (project no. 616803-LEARN2SEE to D.Z). We thank Federica Rosselli for her help in getting started with the neurophysiological recordings and Anna Vasilevskaya for her help in exploring motion processing in deep convolutional neuronal networks.

